# THE STRUCTURE OF DNA IN ANABIOTIC AND MUMMIFIED CELLS

**DOI:** 10.1101/2023.09.07.556722

**Authors:** Yurii F. Krupyanskii, Vladislav Kovalenko, Nataliya Loiko, Eduard Tereshkin, Ksenia Tereshkina, Alexander Popov

## Abstract

Investigations of structural organization of DNA for stressed (increased stress resistance), anabiotic and mummified cells obtained by introducing 4-hexylresorcinol at different concentrations for different stages of cell culture growth using diffraction of synchrotron radiation were done. Experimental studies allow us to conclude that 4-hexylresorcinol is the initiator of the cell transition to the anabiotic and mummified states at stationary state. At pre-stationary state of growth 4-hexylresorcinol initiates a transition of cells to a mummified state but not to anabiotic one. The structure of DNA within a cell in the anabiotic dormant state (practically complete absence of metabolism) and the dormant state (starvation stress) coincide (forms *nanocrystalline* structures). Our data suggest the universality of DNA condensation or the universality of DNA protection by a protein Dps for dormant state, regardless of the type of stress. Mummified state (the complete absence of metabolism) is very different in structure (has no ordering within a cell).

## 1. Introduction

Genomic DNA of Escherichia coli has a surprisingly small size of 1 μm3 in the E. coli nucleoid compared to the equilibrium coil [1] in a dilute solution (500 μm3). This is due to condensation of DNA with a DNA-associated proteins in a cell. Condensation allows DNA to occupy a small volume, but not to lose its functionality. Verma et al. [2] indicates the hierarchical organization of DNA in a nucleoid with three tiers of compaction.

Studying the mechanisms and forms of DNA condensation is important for understanding the survival strategies of bacteria, including pathogens, which will help to address their antibiotic resistance.

Bacteria perceive unfavorable changes in environmental conditions as stress. They try to counteract it by using different strategies and adapting their structure and metabolism [3,4]. And first and foremost, the bacteria’s task is to protect the genetic material of the cell [4].

In actively growing cells, as in other living systems, a dynamic, far from equilibrium order is maintained due to metabolism [5,6]. In the stationary phase, when cells go from a phase of active metabolism to a dormant state, they are forced to resort a physical ways of protecting DNA, such as strong DNA packing, crystallization with nucleoid associated proteins, etc., since biochemical ways of DNA protection becomes impossible.

It turned out that the formation of dormant forms also occurs under the action of another type of stress on the bacterial population - synthetic autoregulator capable to provoke microorganism anabiosis, 4-hexylresorcinol (4HR). [3, 7].

In our previous works, synchrotron radiation diffraction and transmission electron microscopy (TEM) were used to study dormant E. coli cells obtained in the natural cycle of culture development after prolonged starvation. We studied DNA structures in dormant and actively growing cells. We observed new *nanocrystalline, liquid crystalline, and folded nucleosome-like 3D DNA patterns* in dormant cell contrast to growing [8].

Several studies have shown that [3,7] the addition of 4HR at low concentrations (up to 10^−4^ M) to a population of stationary cell *E. coli* led to the formation of forms with increased stress resistance (the ability to maintain viability for a longer time than in control bacteria obtained under the same conditions, but not exposed to 4HR).

A dormant anabiotic (practically complete absence of metabolism) state, resembling the dormant state of cells under starvation stress, appears when 4HR is further added to the cell culture. [3,7].

With a further addition of 4HR introduced into the cell suspension, the number of viable cells drops sharply. Bacteria lose their viability (complete absence of metabolism) at 10^−3^ M of 4HR. [7]. However, these non-viable cells retain their external shape during an observation period of about three years under conditions favorable to autolysis. These cells were named mummified cells or micromummies [4,7] The cell walls of such bacteria thickened [7].

This paper describes the action on the design of condensed DNA of a special kind of stress, different from starvation, the action of synthetic autoregulator capable of provoking microorganism anabiosis, 4-hexylresorcinol (4HR). The aim of the work was to study the structural organization of DNA for stressed (increased stress resistance), anabiotic and mummified cells obtained by introducing 4HR at different concentrations at different stages of cell culture growth using synchrotron radiation diffraction. The obtained data on the DNA structure are compared with the structure of condensed DNA formed as a result of starvation stress.

## 2. Materials and Methods

### 2.1 Synthetic autoregulator of anabiosis

4-hexylresorcinol (4HR), chemical formula - 4-hexylbenzene-1,3-diol, was used as a chemical analogue of the anabiosis factor (Sigma-Aldrich, St Louis, MO). Immediately before the experiment, 4HR working water-ethanol solutions were prepared and added to cell suspensions.

### 2.2 Bacterial strain

The work was performed using *Escherichia coli* BL21-Gold *((E. coli* Gold) bacteria from the collection of FRC CP RAS [9, 10].

### 2.3 Culture Conditions

To obtain pre-stationary and stationary cells, *E. coli* Gold bacteria were cultured in Luria-Bertani (LB) medium (Broth, Miller, VWR, Radnor, PA, USA) supplemented with 150 μg/ml ampicillin for 4.5 and 6 hours, respectively. See [9, 10] for details.

### 2.4 *Preparation of anabiotic and mummified E. coli* Gold cells

*E. coli* Gold cells were grown as described above to stationary (6 h) or pre-stationary (4.5 h) growth phases. Then, an alcoholic solution of 4HR was added to the final concentration: to obtain anabiotic cells, 2·10^−4^ M; mummified cells, 10^−3^ M; lower concentrations of 4HR were also used - 10^−4^ M, 10^−5^ M, 10^−6^ M. Then the cells were incubated in a static mode at a temperature of 23°C for 8 days, and then samples were prepared for structural studies.

### 2.5 Preparation of bacterial samples for X-ray structural studies

Selected aliquots of cell samples were centrifuged for 10 minutes at 10000g. Sediment - cell biomass was collected on a sample holder, which was quickly (within 30 seconds, avoiding sample drying) placed in a working area blown with liquid nitrogen, where it was cooled to 100 K. Then the necessary X-ray diffraction studies were done.

### 2.6 Experiments on germination (restoration to a growing state) of anabiotic and mummified cells

Anabiotic and mummified cells prepared as described above were used as inoculum by adding them to fresh LB nutrient medium. Populations were cultured at 1.5, 14 and 96 hours, taking aliquots for preparation of samples for X-ray diffraction studies.

### 2.7 X-ray studies with the use of synchrotron radiation

X-ray experiments were performed at ESRF synchrotron, station ID23-1 (Grenoble, France) at 100 K. The following beam parameters have been set: 1.6799 Å wavelength, 10 μm width of aperture, time of exposition 5s for each diffraction pattern. Flat detector was PILATUS 6M. Detailed description of scattering experiment one may find in [9,10].

As a sample here, ensembles of bacterial cells were used. Scattering angle (2θ) was the angle formed between incident beam axis and diffraction ring, the corresponding resolution is d.

The lattice spacing can be determined based on the increased scattering intensity I corresponding to the d. Thus, characteristic spacings of ordered structures may be identified and conclusions were drawn from X-ray patterns of bacterial cells about architecture of DNA.

## Results

### 3.1. The effect of 4-hexylresorcinol in various concentrations on the DNA structure (cells in a stationary state)

Fig. 1A shows the scattering curves of synchrotron radiation for cells exposed to 4HR at concentrations of 2·10^−4^ M (blue curve) and 10^−3^ M (red curve), and Figs. 1B shows scattering curves of synchrotron radiation for cells exposed to 4HR at a concentration of 10^−6^ M (green), 10^−5^ M (violet), and 2·10^−4^ M (blue)

**Figure 1.**
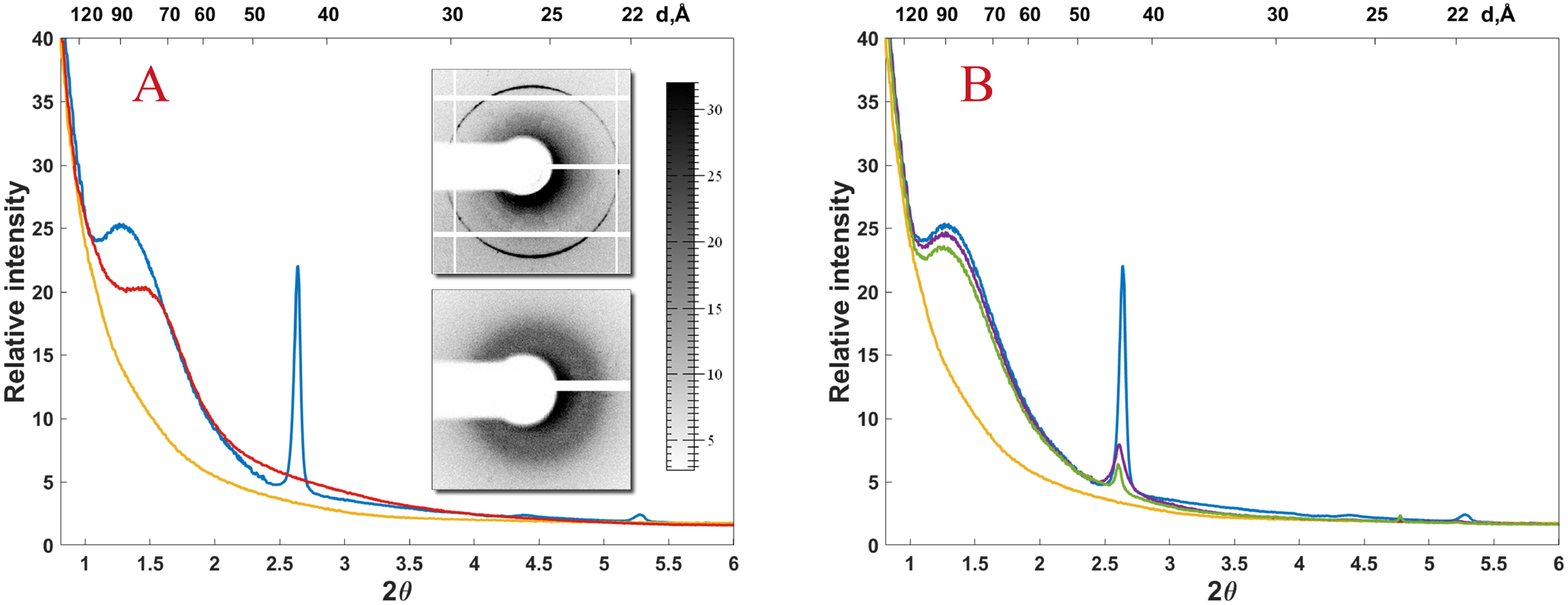
**A** Scattering intensity vs. 2Θ for *E. coli* Gold, stationary growth phase, 4HR was added at a concentration of 2·10^−4^ M, anabiotic state of cell (blue curve) and at a concentration of 10^−3^ M, the mummified state (red curve). The yellow curve (in this figure and throughout) is the scattering intensity for the phase of active growth. The insets show diffraction patterns for the anabiotic and mummified states of cells. **B**. Scatter curves from *E. coli* Gold in the stationary state, exposed to 4HR at a concentration of 10^−6^ M (green), 10^−5^ M (violet), and 2·10^−4^ M (blue curve).

The scattering intensity from *E. coli* cells in the stationary state exposed to 4HR at a concentration of 2·10^−4^ M (blue curve) is identical in shape to the curve obtained from cells under starvation stress [9, 10], see also below Fig.4D. The curve shows peaks corresponding to a resolution of 44 Å and 22 Å and a broad peak with an intensity maximum at a resolution of 88 Å. Such a coincidence of the curves allow to suggest that the organization of DNA in condensed form inside the cells in anabiotic and dormant (starvation stress) state is identical, and that the same Dps protein participates in the formation of protective crystal structures.

With the addition of 4HR up to 10^−3^ M, the effect on cells is significantly different from that observed above. At such a concentration of 4HR micromummies cells appeared [7]. The scattering curve from such cells (red curve in Fig. 1A) has a small broad peak with an intensity maximum corresponding to a resolution of 77 Å. Diffraction peaks corresponding to nanocrystalline structures, like the curves for an anabiotic dormant state on the scattering curves from cells - micromummies are not observed.

As the concentration of 4HR decreases to 10^−5^ M and 10^−6^ M (Fig.1B), the peaks corresponding to 44 Å and 22 Å resolution and the broad peak peaking at 90 Å resolution gradually decrease in intensity.

### 3.2 The effect of 4-hexylresorcinol in various concentrations on the DNA structure (cells in the pre-stationary state)

When 4HR at a concentration of 10^−3^ M is added to *E. coli* Gold cells the scattering curve from these cells (Fig. 2A, blue curve) is almost identical to the scattering curve from cells in the stationary growth phase exposed to 4HR at the same concentration. (Fig.1A). With a decrease in the concentration of 4HR to 2·10^−4^ M, such parallels between the scattering curves of cells in the stationary and pre-stationary state cannot be drawn. The scattering curve from cells in the pre-stationary state does not have Bragg peaks, there is a certain similarity with the smooth scattering curve from growing cells, only the intensity is increased. The curve is even more like the scatter curve from a population grown from anabiotic cells that were in a nutrient medium for 14 hours (Fig. 4C). The identity of the shape of these scattering curves may indicate that the structural response to the effect of 4HR at a concentration of 2·10^−4^ M on the pre-stationary state is like structural response of cells at the beginning of starvation.

**Figure 2.**
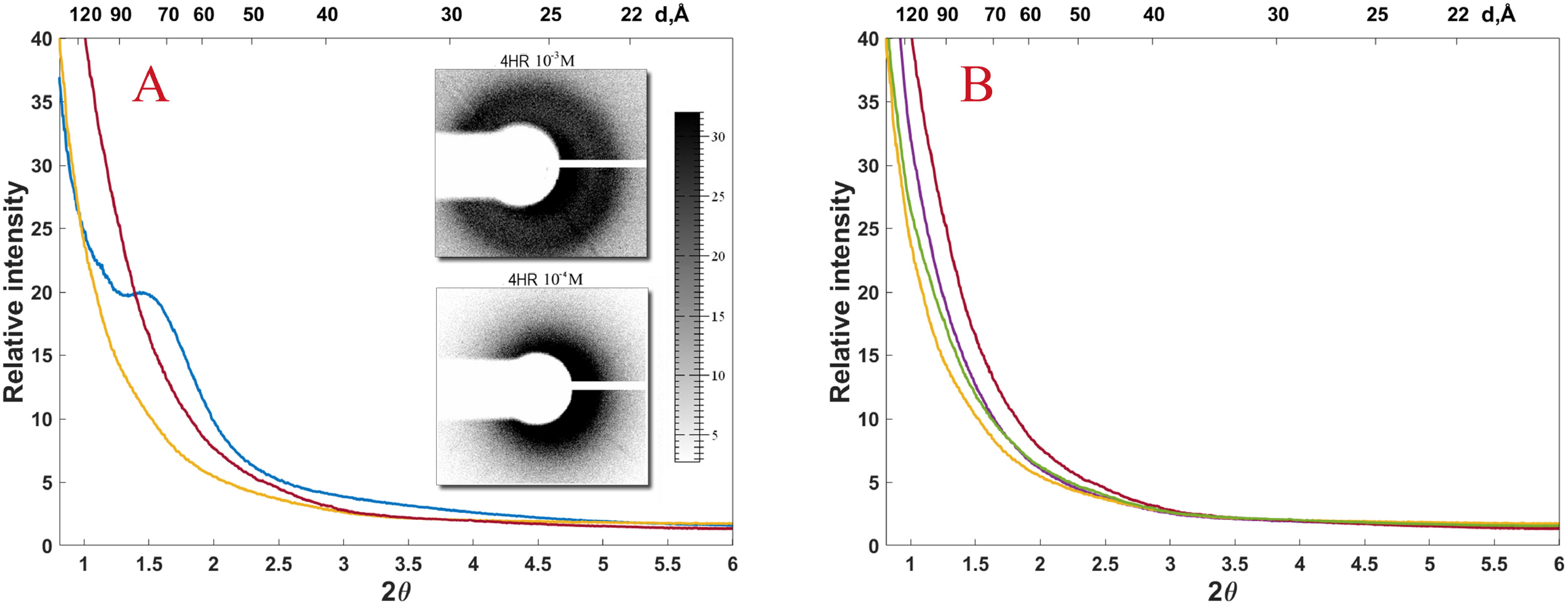
**A**. Scattering intensity vs. 2Θ for *E. coli* Gold, in the pre-stationary phase of development, exposed to 4HR at a concentration of 10^−^3 M (blue curve), 2·10 ^−4^ M, (red curve). **B**. Scattering intensity vs. 2Θ for *E. coli* Gold, in the pre-stationary state, exposed to 4HR at a concentration of 10^−6^ M (green), 10^−5^ M (purple) and 2·10^−4^ M (red curve).

Further, 4HR was introduced into the cell population at concentrations of 10^−6^M, 10^−5^ M and 2·10^−4^M (Fig. 2B). Compared to cells that grow actively, in cells exposed to 4HR at these concentrations, there is no significant change in the structural organization of DNA, the intensity of scattering increase monotonously with a decrease in the scattering angle. (Fig. 2B). The scattering intensity increases monotonically from curve to curve with increasing 4HR concentration (Fig. 2B). This result is like the behavior of scatter curves from a population grown from anabiotic cells that were in a nutrient medium between 1.5 hours (Fig.4B) and 14 hours (Fig.4C).

### 3.3 Time dependence of the DNA structure of cells in the mummified state

The dependence of the scattering intensity for mummified cells (concentration of 4HR is 10^−3^ M, stationary phase) is shown on Fig. 3. In this case, the structural organization of DNA was studied at different periods of exposure to 4HR on cells: 1.5 hours, 9 hours and 100 hours. As shown in Fig. 3, the scattering curves corresponding to different exposure times are nearly identical to each other. This suggests that the structural organization of mummified cells (within the accuracy of measurements) does not change with time.

**Figure 3.**
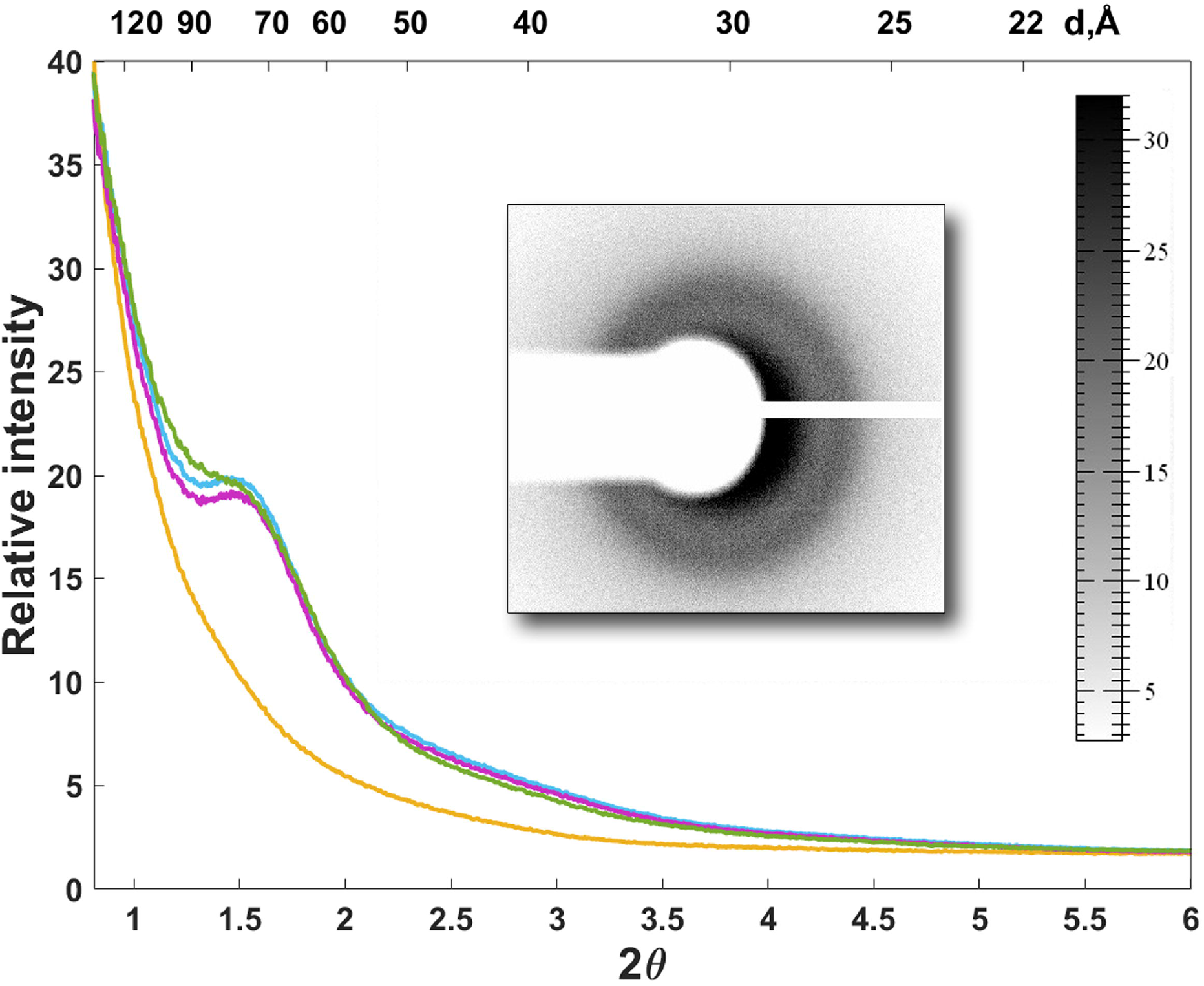
Scattering intensity vs. 2Θ for *E. coli* Gold, stationary state, exposed to 4HR at a concentration of 10^−3^ M for 1.5 hours (blue curve), 9 hours (purple curve), 100 hours (green curve).

### 3.4 Study of the dynamics of activation and germination of anabiotic forms of E. coli

Figure 4A shows the scattering curves from *E. coli* Gold cells stressed with 4HR at 2 x 10^−4^ M and from control population (actively growing cells). Scattering from cells stressed with 4HR contains sharp Bragg peaks at 44 Å and 22 Å and a broad peak with an intensity maximum corresponding to a resolution of 88 Å. For the control population, a smooth monotonic decrease in the scattering intensity is observed with an increase in the angle 2Θ.

**Figure 4.**
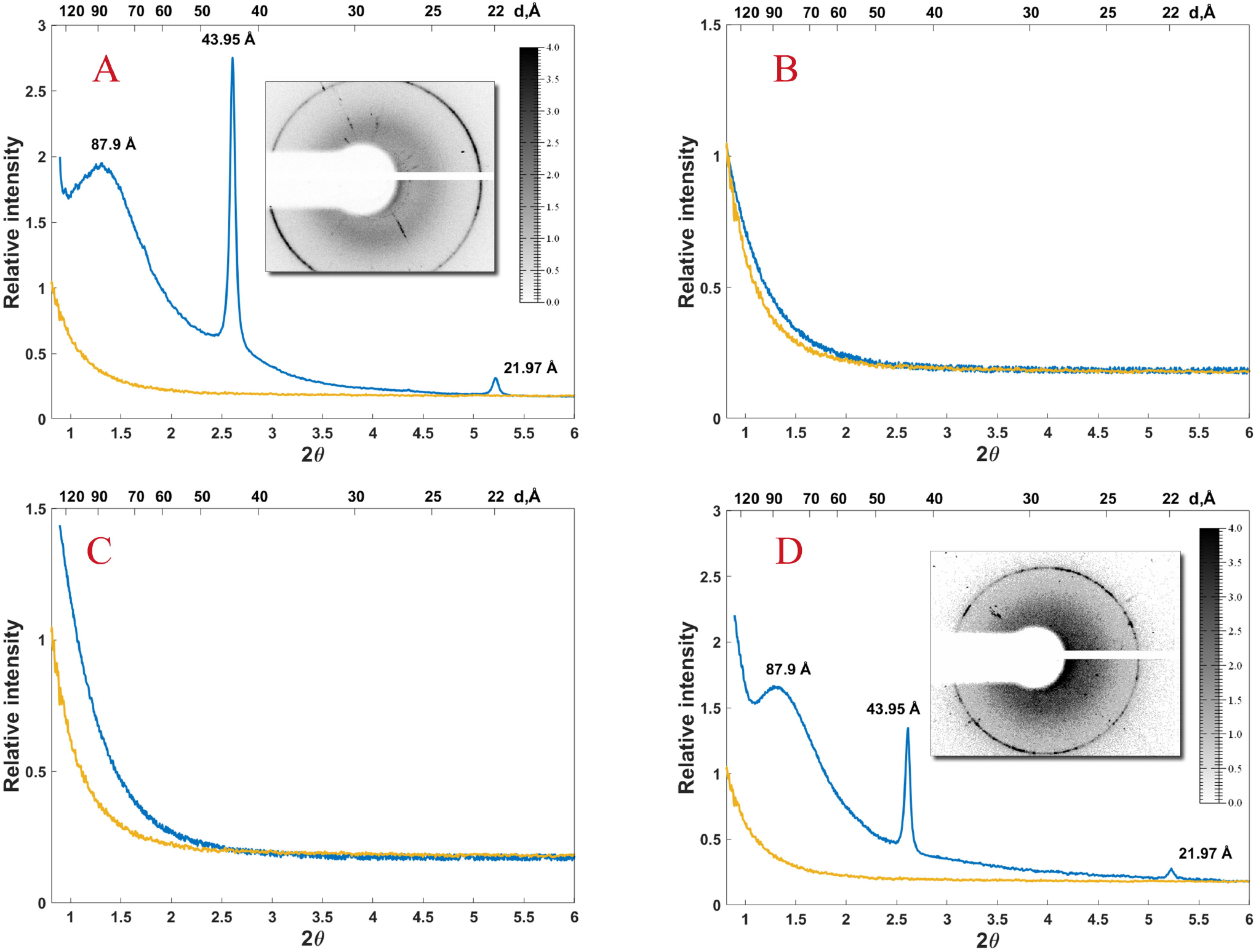
Dynamics of activation and germination of anabiotic forms of *E. coli* Gold, obtained under the action of 4HR (**A)** Yellow curves - growing cells; blue curves - cells exposed to 4HR at a concentration of 2·10^−4^ M (at stationary growth phase); **B** - 1.5 hours in a nutrient medium; **C** - 14 hours in a nutrient medium; **D** - 96 hours in a nutrient medium (starvation stress).

Next, anabiotic *E. coli* cells were placed in a nutrient medium. Fig. 4B shows an almost exact coincidence between the scatter curves from the control population and from anabiotic cells placed in a nutrient medium and cultured in it for an hour and a half. Next, germinated cells were kept for 14 h (Fig. 4C). On Fig. 4C, the difference between the scatter curves from the control population and from the germinated colony begins to appear (scatter intensity is higher). The next step was keeping the germinated cells for 96 hours (Fig. 4D). After 96 hours from the moment the anabiotic cells were placed in the nutrient medium, ordered crystalline formations reappear, the scattering intensity peaks of which are similar in shape, and in position to the peaks obtained from cells corresponding to the stress of exposure to 4HR at a concentration of 2·10^−4^M. This fact allows us to conclude that, according to the diffraction experiment data, the dormant states, both anabiotic and starvation-induced are identical in structure.

## 4. Discussion

During growth, bacterial cells divide into two daughter cells, the genetic material of which is identical to the parent. Four main growth phases of bacterial population can be distinguished. [11]. The initial phase is called the lag phase. Then comes the exponential phase. The population is undergoing exponential growth. Next, bacteria enter stationary phase, in which cells begin to experience starvation stress. If the stress of starvation persists, then the formation of dormant forms occurs. Transition of cells into a dormant state is accompanied by a huge decrease of metabolism. Biochemical protection of DNA in a cell becomes impossible. Cells begin to use physical pathways of protecting DNA, such as strong compaction of DNA, crystallization of DNA with proteins, etc. [6]. It appears that, compared to growing cells, unusual architecture of DNA can be found in dormant cells. In previous works, X-ray diffraction and TEM were used to study dormant *E. coli* cells obtained after prolonged starvation. New structures (*nanocrystalline, liquid crystalline and folded nucleosome-like*) of condensed DNA in dormant cells under starvation stress have indeed been discovered [8].

It was shown in [3,7] that the addition of 4HR at low concentrations (up to 10^−4^ M) to a population of stationary (6 h) *E. coli* Gold cells led to the formation of forms with increased stress resistance, the so-called “stressed” cells (the ability to maintain viability for a longer time than control bacteria obtained under the same conditions, but not exposed to 4HR). An addition of 4HR in the cell culture causes the cells to go into a dormant anabiotic (practically complete absence of metabolism) state, resembling the dormant state of cells at starvation stress. The number of viable cells drops sharply with a further addition of 4HR in the cell suspension. The bacterial cell completely loses its viability (complete absence of metabolism) at concentrations of 4HR above 10^−3^ M. However, these non-viable cells retain their external shape for a period of observation of about three years under conditions conducive to autolysis. These cells were called mummified cells or micromummies [3,7]. The cell membranes of such bacteria thickened.

Fig. 1 (A, C) shows the scattering curves of synchrotron radiation by cells in the stationary growth stage (6 h) exposed from 10^−6^ M up to 10^−3^ M of 4HR. The action of 4HR at a concentration of 10^−6^ M (stationary phase) leads to the appearance of low-intensity peaks corresponding to a resolution of 44 Å and 22 Å and a broad peak corresponding to a resolution of 88 Å (green curve). With increasing 4HR concentration (10^−5^ M), the intensity of the peaks increases (purple curve). At a concentration of 2 10^−4^ M (blue curve in Fig. 1A and Fig. 1B), the cells enter an anabiotic dormant state, very clear intense peaks are observed corresponding to a resolution of 44 Å and 22 Å and a broad peak corresponding to a resolution of 88 Å.

Since exactly the same peaks at the same resolutions we observed earlier for dormant cells (starvation stress), which we associated with the *nanocrystalline* state of condensed DNA, (the TEM [8, 12] data were also used to establish the *nanocrystalline* structure), the coincidence of the scattering curves leads to the idea of the identity of the structure of condensed DNA inside an anabiotic and dormant (starvation stress) cell, and that the same Dps protein is involved in the formation of protective *nanocrystalline* structures.

The low intensity of the peaks in stressed cells compared to anabiotic cells is apparently explained by the smaller size of the crystalline regions in these cells. An increase in the concentration of 4HR increases the size of crystalline regions and number of crystalline regions in cells, reaching a maximum in anabiotic cells at a concentration of 4HR in 2 10^−4^ M.

With a further increase in the concentration of 4HR introduced into the cell suspension, the number of viable cells drops sharply. At concentrations above 10^−3^ M, the action of 4HR causes a mummified state of cell (micromummies) formation [3,7] (Fig. 1A, red curve). The scattering curve for micromummies is completely different from the scattering curve for the anabiotic state of cells. The scattering curve from micromummies has a small broad peak with an intensity maximum corresponding to a resolution of 77 Å. Diffraction peaks corresponding to *nanocrystalline* structures in anabiotic state are not observed on the scattering curves from cells - micromummies. It is not possible to attribute the broad peak at a resolution of 77 Å to any intracellular structure. Apparently, it is associated with scattering on the thickened walls of micromummies.

Next, 4HR was introduced at the pre-stationary state of cell (4.5 h), when the cells had not yet decided on their further growth program, and when the amount of DNA-binding Dps proteins was not yet high. In cells at 4HR in concentrations of 10^−6^ M, 10^−5^ M and 2·10^−4^ M (Fig. 2B), there is no radical change in architecture of DNA compared to actively growing cells, the scattering curves monotonously increase with decreasing scattering angle and do not have characteristic peaks (Figure 2B). However, the intensity of the curves increases with increasing 4HR concentration, which indicates the appearance of new scattering centers, possibly ultrasmall crystallites tens of Å in size, but not yet well-ordered structures inside the cells. These scatter curves (4HR concentration - 10^−6^ M, 10^−5^ M and 2·10^−4^ M) have a certain similarity with the scatter curves from a colony sprouted from anabiotic cells and kept in a nutrient medium from 1.5 hours to 14 hours (Fig. 4B and 4C).

When 4HR at a concentration of 10^−3^ M is added to *E. coli* in the pre-stationary state, the scattering curve from these cells (Fig. 2A) is almost identical to the scattering curve from cells in the stationary growth phase when exposed to 4HR at the same concentration (Fig. 1A). The effect of 4HR on cells at a concentration of 10^−3^ M is very strong and transforms it into micromummy, regardless of whether 4HR is introduced in the pre-stationary (4.5 h) state (the cells have not yet decided on the growth program) or in the stationary state. This concentration of 4HR radically changes the cell growth program and transforms it into micromummy.

Mummified cells completely lose their viability (they completely lack metabolism). These cells retain their external shape for three years under conditions conducive to autolysis [7]. The scattering curves shown in Figure 3 reflect the ability of micromummies not to change shape and, apparently, the internal structure over time. The scattering curves are identical to each other up to 100 hours of exposure (Fig. 3). In the mummified state, intracellular ordered structures are not observed. Apparently, this is the result of the fact that the mummified state is much closer to the state of thermodynamic equilibrium with maximum entropy (or disorder), see [5].

Figure 4 shows the results of studying the dynamics of activation and germination of anabiotic forms of *E. coli*. Scattering from anabiotic cells contains sharp Bragg peaks at 44 Å and 22 Å and a broad peak at 88 Å (Figure 4A). When anabiotic *E. coli* cells are placed in a nutrient medium for 1.5 hours, there is almost an exact coincidence of the scattering curves from the control population and from anabiotic cells in the nutrient medium. The germinated cells were then allowed to stand for 14 h (Fig. 4C). The scattering curve does not change the monotonicity of the character, but the scattering intensity increases noticeably. For cells kept for 96 h, starvation stress begins to show (Fig. 4D). *Nanocrystalline* formations reappear (Fig. 4D), the scattering intensity peaks are almost identical in shape and position to the peaks obtained from anabiotic cells (Fig. 4A). The coincidence of the scattering curves once again indicates the identity of the architecture of DNA within anabiotic and dormant (starvation stress) cells, and that the same Dps protein is involved in the formation of protective *nanocrystalline* structures. Finally, it should be noted here that experimental studies of both the anabiotic dormant state and the mummified state are just at the beginning.

## Acknowledgments

Authors are grateful to ESRF ID23-1 (Grenoble, France) for the possibility of carrying out X-ray experiments. The results were processed at JSCC RAS.

